# *De novo* assembly, characterization, functional annotation and expression patterns of the black tiger shrimp (*Penaeus monodon*) transcriptome

**DOI:** 10.1101/280420

**Authors:** Roger Huerlimann, Nicholas M Wade, Lavinia Gordon, Juan D Montenegro, Jake Goodall, Sean McWilliam, Matthew Tinning, Kirby Siemering, Erika Giardina, Dallas Donovan, Melony J Sellars, Jeff A Cowley, Kelly Condon, Greg J Coman, Mehar S Khatkar, Herman W Raadsma, Gregory Maes, Kyall R Zenger, Dean R Jerry

## Abstract

The black tiger shrimp (*Penaeus monodon*) remains the second most widely cultured shrimp species globally. However, issues with disease and domestication have seen production levels stagnate over the past two decades. To help identify innovative solutions needed to resolve bottlenecks hampering the culture of this species, it is important to generate genetic and genomic resources. Towards this aim, we have produced the most complete publicly available *P. monodon* transcriptome database to date. The assembly was carried out in multiple assemblers using 2×125 bp HiSeq data from PolyA selected, ribo-depleted RNA extracted from nine adult tissues and eight early life-history stages. In total, approximately 700 million high-quality sequence reads were obtained and assembled into 236,388 clusters. These were then further segregated into 99,203 adult tissue specific clusters, and 58,678 early life-history stage specific clusters. The final transcriptome had a high TransRate score of 0.37, with 88% of all reads successfully mapping back to the transcriptome. BUSCO statistics showed the assembly to be highly complete with low fragmentation, few genes missing, but higher redundancy or transcript duplication (Complete: 98.2% (Duplicated: 51.3%), Fragmented: 0.8%, Missing: 1.0%), and to greatly exceed the completeness of existing *P. monodon* transcriptomes. While annotation rates were low (approximately 30%), as is typical for a non-model organisms, annotated transcript clusters were successfully mapped to several hundred functional KEGG pathways. To help address the lack of annotation, transcripts were clustered into groups within tissues and early life-history stages, providing initial evidence for their roles in specific tissue functions, or developmental transitions. Additionally, transcripts of shrimp viruses previously not known to occur in Australia were also discovered. We expect the transcriptome to provide an essential resource to investigate the molecular basis of commercially relevant-significant traits in *P. monodon* and other shrimp species.

## Introduction

The black tiger shrimp *Penaeus monodon* belongs to the family Penaeidae and is the second most widely farmed shrimp species globally^1^. However, disease and limited progress in domestication and selective breeding of *P. monodon* continue to hamper further expansion of the industry^2^. Modern genomic technologies have significant potential to advance selective breeding programs; however, they require complete, well annotated tissue-specific transcriptomic and genomic datasets. In addition to assisting in genome assembly and creating linkage maps^3^, a complete transcriptome provides a potential resource for differential gene-expression studies^4^), genome annotation^5^, single nucleotide polymorphism discovery^6^ and genome scaffolding^7^.

While genomic resources for Penaeid shrimp are increasing, they remain limited for many species, including *P. monodon*. Previous research has focussed on hepatopancreas, ovary, heart, muscle and eyestalk tissues^8,9^, in male and female gonads^10^, and in response to infection with *Vibrio* bacterial species capable of inducing acute hepatopancreatic necrosis disease^11^. In addition to such differential gene-expression studies, genomic data from next generation sequencing (NGS) methods has expanded greatly in recent years, particularly in the study of Pacific white shrimp (*Litopenaeus vannamei*)^3,6,12–23^. Moreover, a transcriptome based on eight tissues was assembled for the less well studied banana shrimp *Fenneropenaeus merguiensis*^24^, and genes involved in early embryonic specification have been studied in *Marsupenaeus japonicus*^25^. Transcriptomics has also been applied to *Penaeus merguiensis*^26–28^ and the Chinese white shrimp *Fenneropenaeus chinensis*^29,30^ to investigate aspects of tissue-specific expression, stress tolerance and viral infection. Despite these advances, a comprehensive transcriptome from diverse tissue types and early life-history stages of *P. monodon* remains unavailable.

In order to address this deficiency, we report a highly complete transcriptome for *P. monodon* that can be used as a broad basis for future genomics research. To this effect, we sequenced three replicates each from nine different tissues types (eyestalk, stomach, female gonad, male gonad, gill, haemolymph, hepatopancreas, lymphoid organ and tail muscle) and one pooled replicate each from four larval stages (embryo, nauplii, zoea, and mysis) and four post-larval stages ranging from days 1, 4, 10 and 15. Additionally, transcript expression profiles unique to each type and stage were determined, as well as identifying putative long non-coding RNA and transcripts originating from viruses.

## Material and Methods

### Sample taking and RNA extraction

Tissues of *P. monodon* broodstock were collected from multiple individuals, immediately snap frozen on dry ice, and stored at −80°C until extraction (Table 1). All tissues except lymphoid organs were collected from wild broodstock caught off coastal waters near the border between the Northern Territory and Western Australia provided, which were provided by a commercial hatchery at Flying Fish Point, North Queensland, Australia. Lymphoid organ tissue was collected from wild prawns caught off the East Coast of Queensland. Larval and post-larval stages were collected from the same hatchery in pools of approximately 400 individuals per life stage, after four hours of starvation, and preserved in RNAlater (Thermo Fisher Scientific). All tissues and early life-history stages were sub-sampled in an RNase-free laboratory and total RNA was extracted using an RNeasy Universal extraction kit (QIAGEN) following manufacturer’s instructions. RNA quantity and quality was estimated using a Nanodrop UV spectrophotometer (Thermo Fisher Scientific), and purity was further assessed using an Agilent Bioanalyzer (Agilent Technologies).RNA was selected from individual sample replicates based on Nanodrop spectra, RNA concentration, and Agilent Bioanalyzer traces (Table 1), in preference to using comparative tissues from the same individuals.

**Table 1.** List of shrimp tissue types and early life-history stages used for transcriptome sequencing. PL = post-larval stages 1 (PL1), 4 (PL4), 10 (PL10), 15 (PL15)

| Shrimp ID | Sex | Tissue | Number of paired-end reads |
| --- | --- | --- | --- |
| PM_F_08 | Female | Eyestalk | 18,984,152 |
|  |  | Gill | 19,971,115 |
|  |  | Hepatopancreas | 18,831,682 |
| PM_F_02 | Female | Female Gonad | 21,338,933 |
|  |  | Haemolymph | 20,105,399 |
|  |  | Muscle | 20,361,299 |
|  |  | Stomach | 13,470,106 |
| PM_F_04 | Female | Female Gonad | 20,255,448 |
|  |  | Gill | 21,362,076 |
|  |  | Haemolymph | 20,247,206 |
|  |  | Stomach | 21,461,589 |
| PM_F_03 | Female | Female Gonad | 20,759,890 |
| PM_M_02 | Male | Eyestalk | 21,076,111 |
|  |  | Hepatopancreas | 19,029,973 |
|  |  | Male Gonad | 20,669,419 |
|  |  | Muscle | 20,129,858 |
| PM_M_04 | Male | Eyestalk | 22,250,295 |
|  |  | Gill | 20,396,956 |
|  |  | Haemolymph | 21,637,767 |
|  |  | Hepatopancreas | 20,854,492 |
|  |  | Male Gonad | 20,600,256 |
|  |  | Muscle | 22,464,431 |
|  |  | Stomach | 16,444,377 |
| PM_M_06 | Male | Male Gonad | 19,800,274 |
| PM_M_C2 | Male | Lymphoid Organ | 19,873,753 |
| PM_M_C3 | Male | Lymphoid Organ | 20,480,178 |
| PM_F_C1 | Female | Lymphoid Organ | 20,372,862 |
| Pool_E |  | Embryo | 19,745,313 |
| Pool_N |  | Nauplii | 18,310,089 |
| Pool_Z |  | Zoea | 19,528,689 |
| Pool_M |  | Mysis | 19,744,563 |
| Pool_PL1 |  | PL1 | 19,815,103 |
| Pool_PL4 |  | PL4 | 18,680,555 |
| Pool_PL10 |  | PL10 | 18,773,667 |
| Pool_PL15 |  | PL15 | 19,661,826 |

### Illumina library preparation and sequencing

Library preparation and sequencing was carried out at the Australian Genome Research Facility (AGRF). Upon arrival at the sequencing facility, the quality of the samples was checked using a Bioanalyzer RNA 6000 nano reagent kit (Agilent) and libraries were prepared using the TruSeq Stranded mRNA Library Preparation Kit (Illumina) according to established protocols. Final libraries were again checked using Tapestation DNA 1000 TapeScreen Assay (Agilent). Cluster generation was performed on a cBot with HiSeq PE Cluster Kit v4 - cBot and sequencing was done on a HiSeq 2500 using a HiSeq SBS Kit. The Hiseq 2500 was operating with HiSeq Control Software v2.2.68 and base-calling was performed with RTA v1.18.66.3.Samples in the second sequencing run were pooled and split across two lanes to reduce sequencing bias (Table 1).

### Sequence quality control, assembly and annotation

Raw sequence data was quality checked using FastQC^31^ v0.11.5, and assembled loosely following the Oyster River Protocol for Transcriptome Assembly^32^. In brief, all sequences were collectively error-corrected using RCorrector^33^ V3. Samples were then assembled in Trinity^34^ V2.3.2; grouped by individual shrimps, i.e. all tissues from a specific shrimp were assembled together. Reads were trimmed harshly for adapters and softly for Phred score <2 using Trimmomatic^35^ V0.32; and then normalized *in silico* within Trinity. The normalized forward and reverse reads produced by Trinity were then used in BinPacker^36^ V1.0, IDBA-Tran^37^ V 1.1.1 using K20, K30, K40, K50 and K60; and Bridger^38^ version 2014-12-01. All resulting transcriptomes were concatenated and merged using Evidential Gene^39^, followed by clustering using Transfuse V0.5.0 (https://github.com/cboursnell/transfuse) using a similarity value of 0.98. Lastly, contigs <300 bp were removed to produce the final transcriptome. The quality of the final assembly was assessed using TransRate^40^ V1.0.1, and BUSCO^41^ V2 using the arthropoda_odb9 database^42^. Sequences were annotated in Blast2Go^43^ using the SWISS-PROT database^44^ (accessed 17/03/2017), and separately using the arthropod and viral subsections of the GenBank nr database (accessed 06/06/2017).

### Identification of long non-coding RNAs

FEELnc^45^ was used for the identification of long non-coding RNAs. The coding transcripts training set was constructed from the 1,047 complete universal single copy orthologous genes found with BUSCO v2.0 (database arthropoda_odb9^42^). The mode “shuffle” was used to generate a training set of lncRNA from the debris of the known coding RNA transcripts.

### Mapping and differential gene expression analysis

Before mapping, error-corrected raw sequence reads were trimmed using the same parameters as before, but without palindrome trimming used by Trinity. Sequence reads were mapped using Bowtie2^46^ V2.2.8, and read counts were calculated using Corset^47^ V1.0.6. Differential gene expression was analyzed using DESeq2^48^ V1.16.1 in RStudio^49^ V3.4.1.

To reduce the number of sequences for KEGG analysis, the longest contig per cluster was chosen from the combined tissue type and early life-history stage data. The KEGG Automatic Annotation Server (KAAS, http://www.genome.jp/tools/kaas/) was used to generate KEGG pathway maps for each contig using BLAST with the single-directional best hit (SBH) method. All scripts will be deposited on GitHub upon acceptance.

### Statistical analyses

For data analysis, the top 2,000 variably expressed genes across the nine tissue types and the top 500 variably expressed genes across the four larval and four post-larval stages were visualized in a principal component analysis and heatmap using variance-stabilizing transformed read-count data from DESeq2. The gene level dendrograms in the heatmap were created using Pearson’s correlation for both the tissue type larval/post-larval stages. Euclidean distance was used to cluster tissue types. All statistical analyses were performed in RStudio. Detailed information on the analyses can be found on GitHub upon acceptance.

## Results

### Sequence read data and code availability

In total, nine tissues were sequenced in biological triplicates, as well as pools of eight early life-history stages, resulting in an average of 19.9 M ± 1.6 M (mean ± SD) read pairs per sample and 697 M reads in total (Table 1). After quality trimming, 99.5% ± 0.6% (mean ± SD) of reads were retained, indicating a high quality data set (>90% reads with ≥Q30). All read data are available on GenBank through the project ID PRJNA421400.

### Transcriptome assembly and quality control

The initial combined outputs of all four assemblers comprised of 6,113,055 contigs, which were reduced to 462,772 contigs after filtering with Evidential Gene and combining both “okay” and “alternative” contigs. After clustering with Transfuse, the final assembly consisted of 236,388 transcripts with an assembly size of 226 Mb.These, together with transcript annotations, are available on GenBank. The final transcriptome had a high TransRate score of 0.37, with 88% of all reads successfully mapping back to the transcriptome, and only 3.2% of bases being uncovered. Based on BUSCO, the transcriptome was highly complete with 98% of arthropod ortholog genes being present, and few fragmented or missing genes; however, 51% of the contigs were duplicated/redundant (C:98.2%[S:46.9%,D:51.3%],F:0.8%,M:1.0%,n:1066).

### Annotation and gene ontology mapping

Annotation against the SwissProt database using BLASTx resulted in 47,871 successfully annotated contigs. Of these, 46,977 were successfully GO mapped, of which 41,069 were completely annotated. The top-hit species distribution was dominated by *Homo sapiens* with over 10,000 hits, followed by *Drosophila melanogaster* with just over 8,000 hits; no shrimp species made it into the list (Fig. 1). GO terms for biological processes, molecular function and cellular components were all highly represented in annotated genes (Fig. 2).

**Figure 1.**
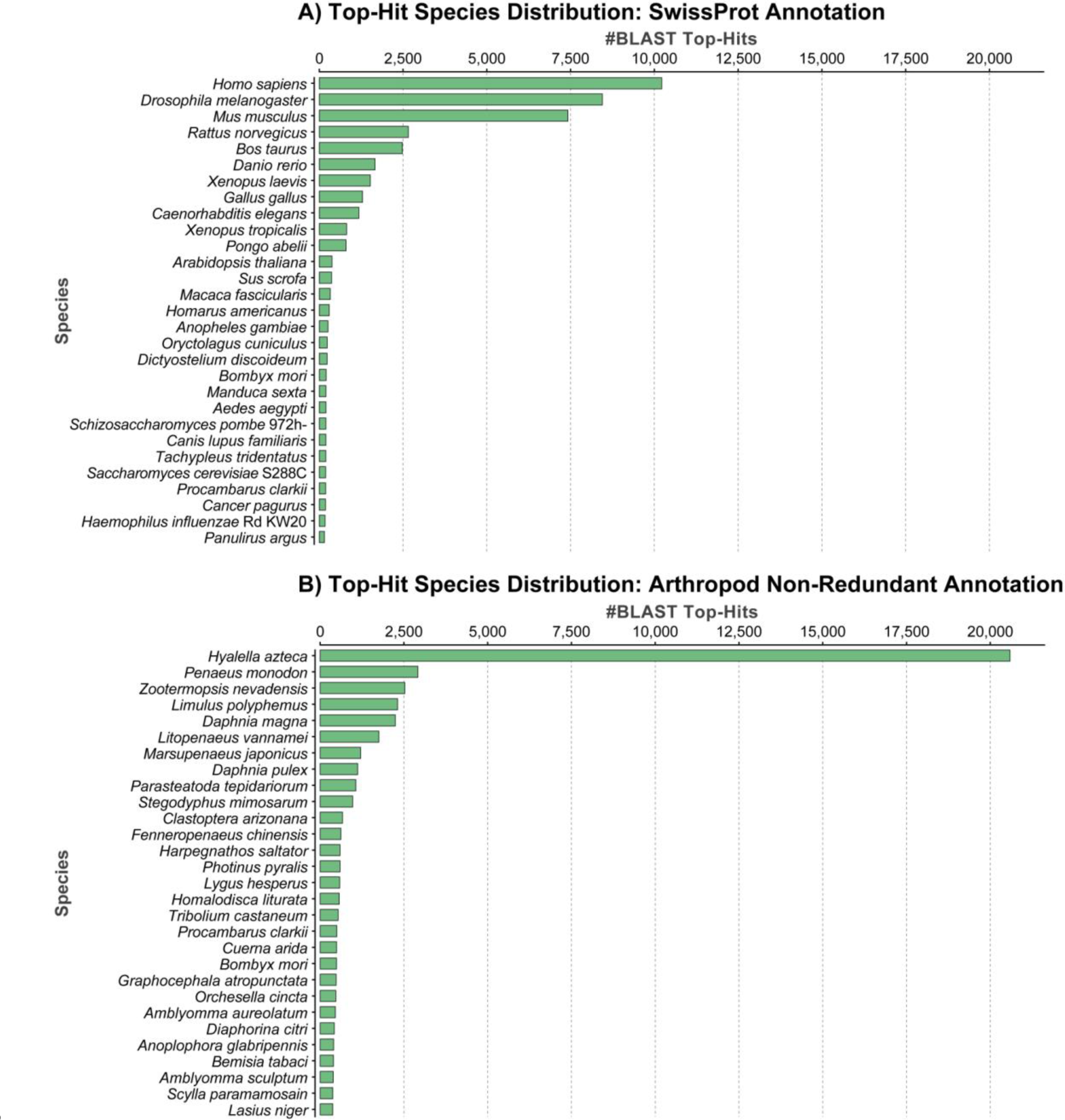
Species distribution of successfully annotated sequences across the top 29 species using the SwissProt (A) and arthropod subsection of the non-redundant (B) database.

The annotation against the non-redundant Arthropod (nrA) database using BLASTx resulted in 62,679 successfully annotated contigs, of which 48,456 had a successful GO mapping, and of which 25,201 were completely annotated. The top-hit species distribution was dominated by the freshwater amphipod *Hyalella azteca* with over 20,000 hits, followed by *P. monodon* with just over 2,500 hits (Fig. 1). Other penaeid shrimp species included *Litopenaeus vannamei, Marsupenaeus japonicus* and *Fenneropenaeus chinensis*, which were the sixth, seventh and twelfth most highly represented species respectively.

Detailed information on the annotations can be found in Supplementary Table 1.

**Figure 2.**
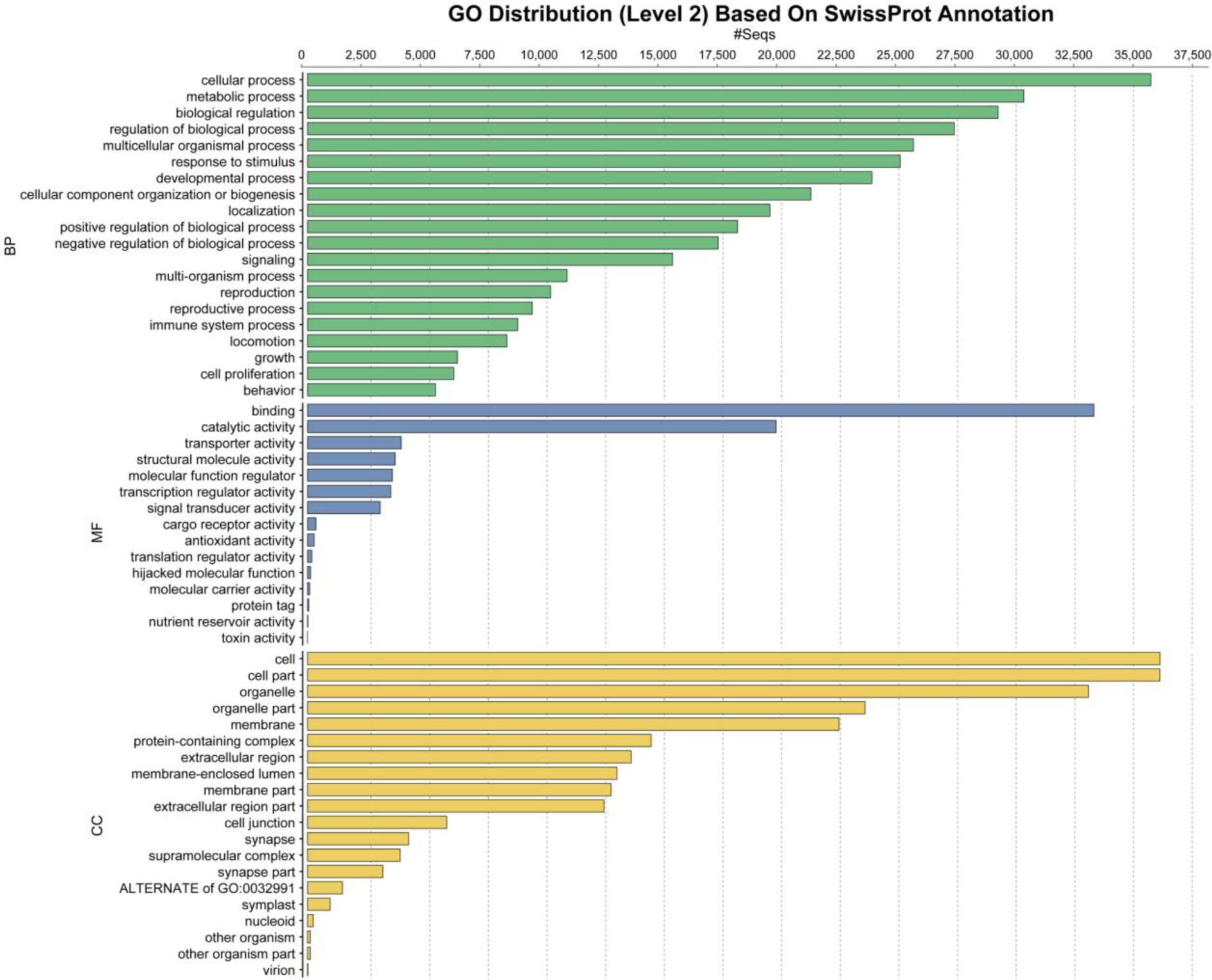
Distribution of sequence annotations based on the SWISS-PROT database across the top 20 GO terms at level 2. Determined across the entire dataset for Biological Process (BP, green), Molecular Function (MF, blue), and Cellular Component (CC, yellow).

### Sequence read mapping and differential gene expression analysis

Using Bowtie2, 67.4% ± 4.8% (mean ± SD) of the paired reads successfully mapped to the transcriptome. Using corset for read counting and additional clustering, the initial 236,388 contigs were placed into 99,203 transcript clusters for the nine tissue types and 58,678 transcript clusters for the eight early life-history stages (larval and post-larval stage). A total of 176,966 contigs were used in the clustering of tissues and larvae, with 113,435 shared contigs, 8,188 contigs unique to larvae and 55,343 contigs unique to adult tissues.

Different tissue types expressed between 9,939 and 12,255 transcript clusters (defined as > 50 normalized read counts per cluster), and between 17 and 316 unique sets of transcript clusters (defined as a cluster with > 10 normalized read counts and < 10 normalized read counts in all other tissue types) (Table 3). The ability to annotate transcript clusters varied across tissue types (63.0% to 85.9%). In terms of unique tissue specific transcript clusters, hepatopancreas contained the largest number (316), followed by female gonad (161) and gill (153). Annotation rates of these unique tissue-specific clusters were markedly lower (12.5% to 66.8%) than with clusters shared across all tissue types (82.5% and 85.9%)

**Table 3.** Numbers of transcript clusters and cluster annotation rates across transcriptomes determined for the nine adult *P. monodon* tissue types analysed. Total numbers of expressed clusters (>50 normalized read counts), uniquely expressed clusters (normalized read count of >10 in a specific tissue, while having <10 read counts in all other tissues) and constitutively expressed (> 50 normalized read counts in all) clusters within all tissues in this study, and their relative annotation statistics. Numbers represent clusters across all three respective tissue replicates. SP = SWISS-PROT database, nrA = non-redundant Arthropod database.

| Tissue type | Total expressed clusters |  | Uniquely expressed clusters |  |
| --- | --- | --- | --- | --- |
|  | Number | % Annotated (SP/nrA) | Number | % Annotated (SP/nrA) |
| Eyestalk | 11,173 | 67.3 / 72.8 | 31 | 29.0 / 48.4 |
| Female Gonad | 9,941 | 74.3 / 79.7 | 161 | 37.3 / 45.3 |
| Gill | 12,255 | 63.7 / 69.8 | 153 | 30.7 / 39.2 |
| Haemolymph | 10,577 | 66.1 / 71.4 | 17 | 23.5 / 29.4 |
| Hepatopancreas | 12,169 | 67.7 / 73.9 | 316 | 49.7 / 66.8 |
| Lymphoid Organ | 11,923 | 63.0 / 68.5 | 24 | 54.2 / 66.7 |
| Male Gonad | 10,387 | 71.9 / 77.5 | 71 | 32.4 / 42.3 |
| Muscle | 11,405 | 66.9 / 72.4 | 77 | 33.8 / 48.1 |
| Stomach | 9,939 | 68.6 / 73.7 | 24 | 12.5 / 33.3 |
| Constitutive | 4,300 | 82.5 / 85.9 | - | - |

A principal component analysis (PCA) of the top 1,000 differentially expressed transcripts across the nine adult tissue types showed strong clustering for most tissue replicates, with the exception of stomach and eyestalk (Fig. 3A).Haemolymph, female gonad and muscle formed distinct clusters separated from other tissues, while eyestalk, gill, haemolymph, lymphoid organ, male gonad and stomach tissues were much more closely associated and showed less distinct clustering (Fig. 3A). A PCA of the top 500 differentially expressed transcripts across the eight early life-history stages showed a strong separation within PC1, with embryo and nauplii segregating substantially from the other early life-history larval stages (Fig. 3B). PC1 explained an extraordinary 77% of the variance in transcript clusters expressed across the different discrete larval stages, which appears to be strongly associated with larval development leading from embryo to post-larval stages.

**Figure 3.**
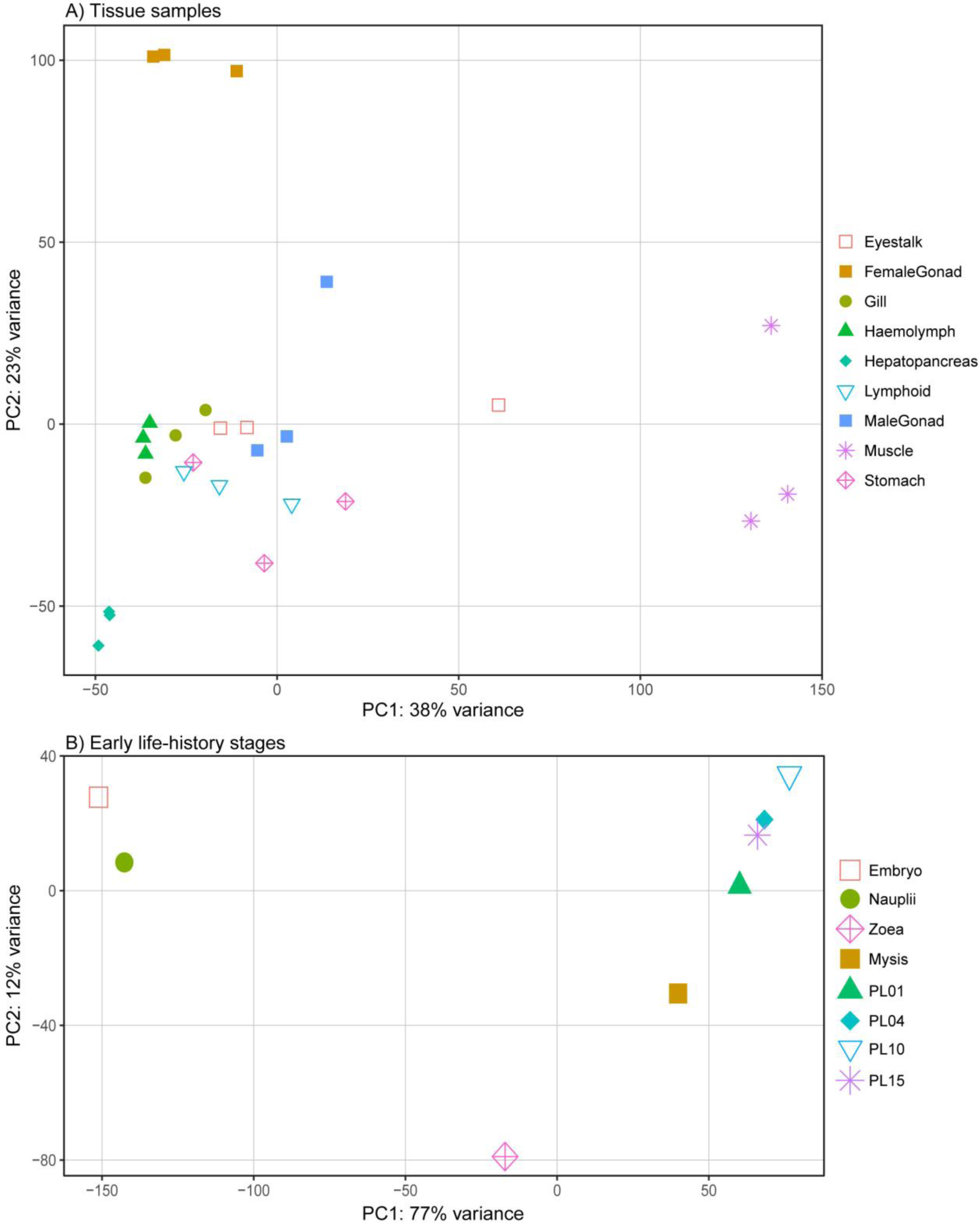
Principal component analysis showing the top most highly differentially expressed transcripts of A) nine tissue types (top 1,000) and B) eight early life-history stages (top 500). PC = principal component, PL = post-larvae

The top 2,000 most variably expressed transcript clusters across all nine tissue types clustered into nine distinct groups using Pearson’s correlation (Fig. 4). These groups aligned broadly with expression patterns identified to be unique to each tissues type. For example, group two comprised 208 clusters highly expressed in female gonad, which were mostly successfully annotated (81.8%) using the nrA database.

**Figure 4.**
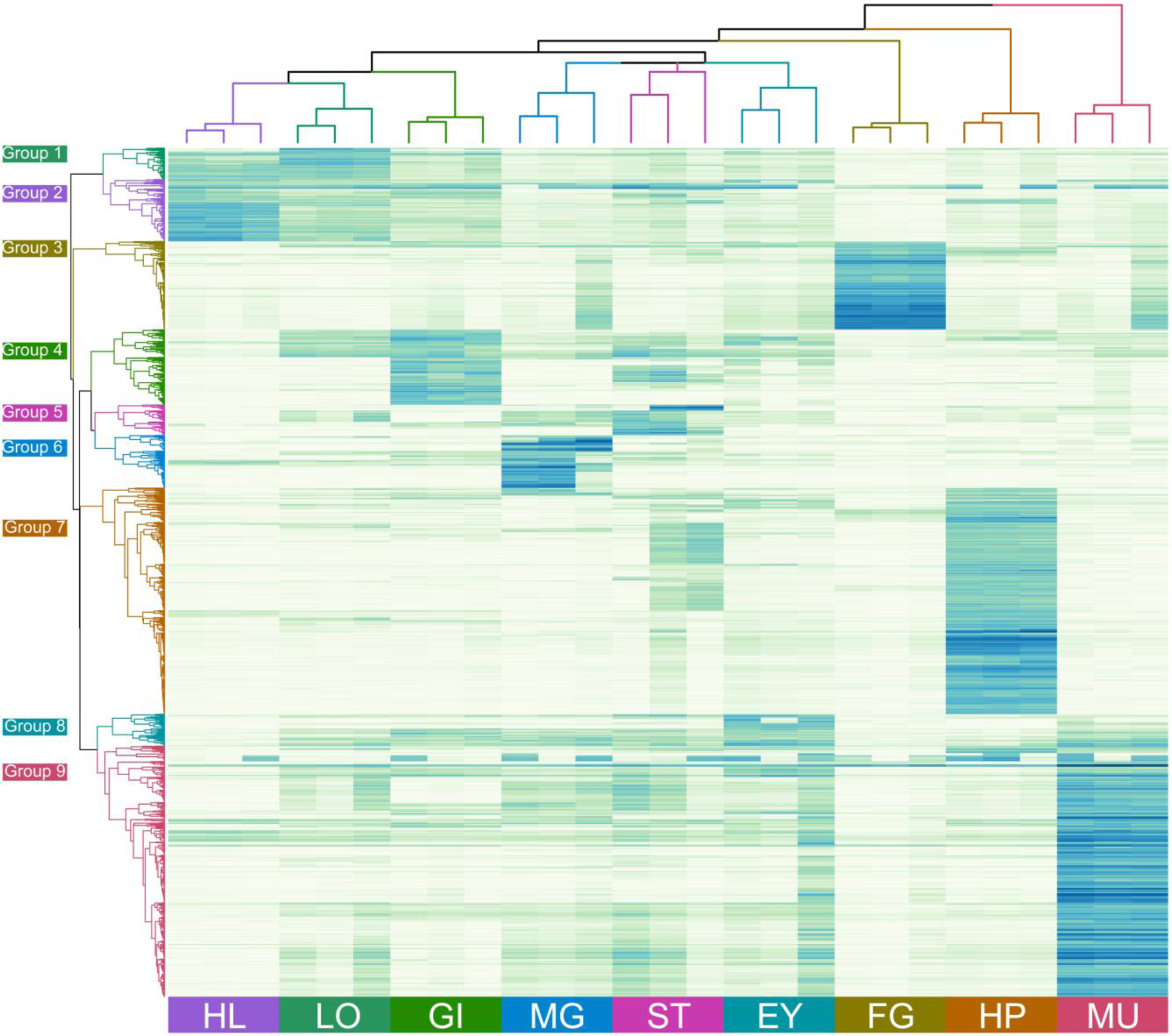
Heatmap and hierarchical grouping of the top 2,000 differentially expressed genes in the nine different tissue types. Gene expression patterns (rows) were grouped into nine expression groups based on Pearson’s correlation and the three replicates of each tissue type (columns) into nine tissue groups based on Euclidean distance. EY – eyestalk; FG – female gonad; GI – gill; HL – hemolymph; HP – hepatopancreas; LO – lymphoid organ; MG – male gonad; MU – muscle; ST – stomach.

Annotated transcripts included farnesoic acid O-methyltransferase (FAmET), phosphoenolpyruvate carboxykinase (PEPCK), glutathione peroxidase (GPx) and nasrat. Transcripts in each cluster and their annotation are detailed in Supplementary Table 2. Group four consisted of clusters expressed mainly in male gonad that were annotated relatively poorly (38.7%) with many (35.5%) not expressed in the early life-history stages (Table 4). Group nine was the largest and comprised 591 clusters that were mostly annotated (86.0%) and expressed predominantly in muscle tissue. Group seven consisted of 533 clusters that were also mostly annotated (85.7%) and expressed predominantly in hepatopancreatic tissue. Except for male gonad, most clusters expressed in adult tissue types were also expressed in the early life-history stages.

**Table 4.** Groupings of the top 2,000 highly variably expressed transcript clusters among all nine adult tissue types based on Pearson’s correlation. This includes annotation success and tissue type where each group was predominantly expressed, and the percent of clusters in each group found in adult tissue types but not in the larval stages examined.

| Groups | Predominant tissue type expression site | Number of clusters | % Annotated (SP/nrA) | % in adult but not larval tissues |
| --- | --- | --- | --- | --- |
| 1 | Lymphoid Organ | 81 | 64.2% / 76.5% | 0.0% |
| 2 | Haemolymph | 139 | 63.3% / 84.9% | 1.4% |
| 3 | Female Gonad | 208 | 55.3% / 81.7% | 6.7% |
| 4 | Gill | 177 | 53.1% / 66.1% | 3.4% |
| 5 | Stomach | 72 | 62.5% / 68.1% | 8.3% |
| 6 | Male Gonad | 124 | 29.0% / 38.7% | 35.5% |
| 7 | Hepatopancreas | 533 | 66.6% / 85.7% | 0.8% |
| 8 | Eyestalk | 75 | 66.7% / 73.3% | 1.3% |
| 9 | Muscle | 591 | 75.1% / 86.0% | 1.5% |
| all | - | 2000 | 64.0% / 84.6% | 4.3% |

SP = SWISS-PROT database, nrA = non-redundant Arthropod database

The same top 500 most variably expressed transcript clusters in the different larval and post-larval stages used for the PCA broadly clustered into nine distinct groups based on Pearson’s correlation (Fig. 5). Irrespective of the annotation success, the analysis identified transcript clusters that shared similar expression patterns across developmental stages. Embryos and nauplii expressed a set of genes that were not expressed during any other developmental stage (groups 7 and 8). Of the 140 genes expressed exclusively within the embryo and nauplii stages (group 8), only 24.3% and 37.1%, respectively, were annotated successfully using the SWISS-PROT or nrA databases (Table 5). Of the transcript clusters that were annotated, 13 encoded orthologs of the neurotrophic factor *spaetzle* and another 13 encoded orthologs of cuticular proteins. Transcripts in each cluster and their annotation are detailed in Supplementary Table 3. Two large clusters of genes were expressed from zoea throughout each subsequent stage (group 1), or from mysis throughout each subsequent stage (group 4). A high percentage (61.2% and 83.1%) of transcripts in these two clusters was annotated. Since each larval stage was sequenced as a pool of individuals, differential gene expression (DGE) analysis could not be performed.

**Table 5.**
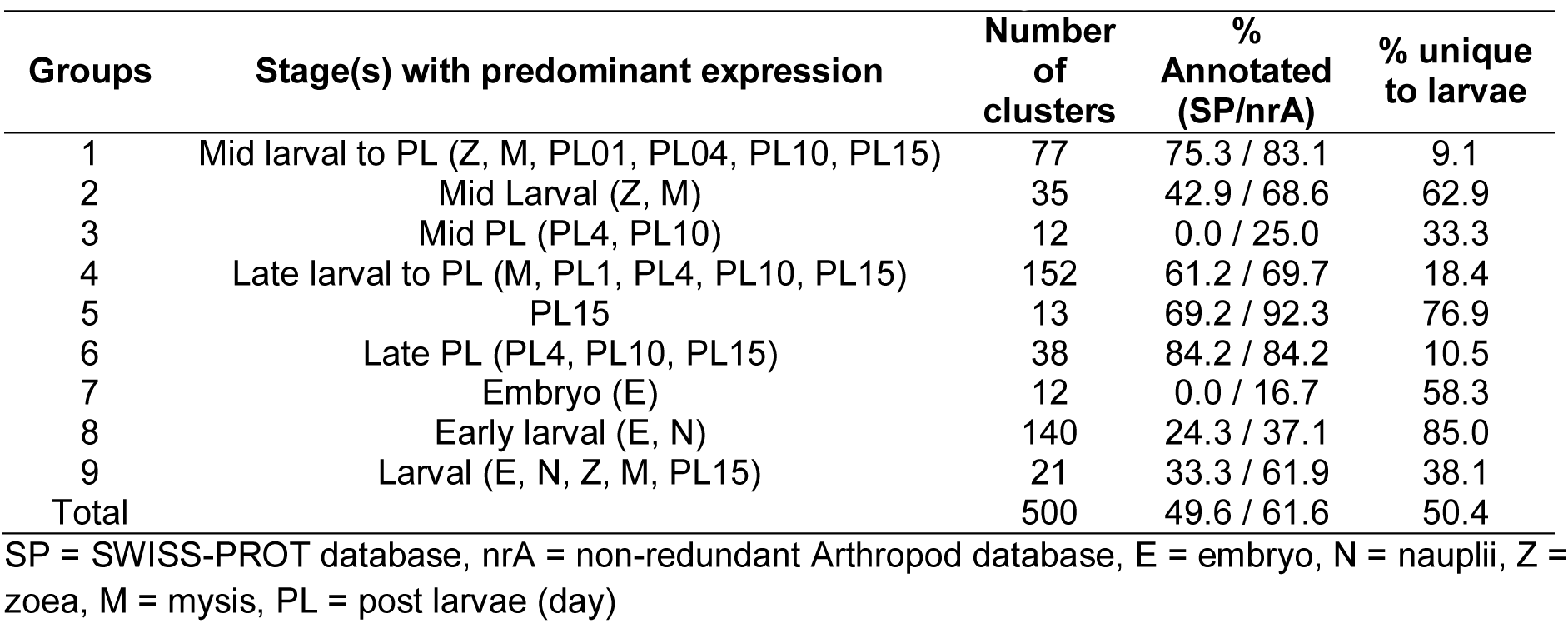
Groupings of the top 500 highly variably expressed transcript clusters among the four larval and four post-larval stages based on Pearson’s correlation. This includes annotation success, stages in which transcript groups were predominantly expressed and the percent of clusters in each group found in larval stages, but not in the adult tissue types examined.

| Groups | Stage(s) with predominant expression | Number of clusters | % Annotated (SP/nrA) | % unique to larvae |
| --- | --- | --- | --- | --- |
| 1 | Mid larval to PL (Z, M, PL01, PL04, PL10, PL15) | 77 | 75.3 / 83.1 | 9.1 |
| 2 | Mid Larval (Z, M) | 35 | 42.9 / 68.6 | 62.9 |
| 3 | Mid PL (PL4, PL10) | 12 | 0.0 / 25.0 | 33.3 |
| 4 | Late larval to PL (M, PL1, PL4, PL10, PL15) | 152 | 61.2 / 69.7 | 18.4 |
| 5 | PL15 | 13 | 69.2 / 92.3 | 76.9 |
| 6 | Late PL (PL4, PL10, PL15) | 38 | 84.2 / 84.2 | 10.5 |
| 7 | Embryo (E) | 12 | 0.0 / 16.7 | 58.3 |
| 8 | Early larval (E, N) | 140 | 24.3 / 37.1 | 85.0 |
| 9 | Larval (E, N, Z, M, PL15) | 21 | 33.3 / 61.9 | 38.1 |
| Total |  | 500 | 49.6 / 61.6 | 50.4 |

SP = SWISS-PROT database, nrA = non-redundant Arthropod database, E = embryo, N = nauplii, Z = zoea, M = mysis, PL = post larvae (day)

**Fig. 5.**
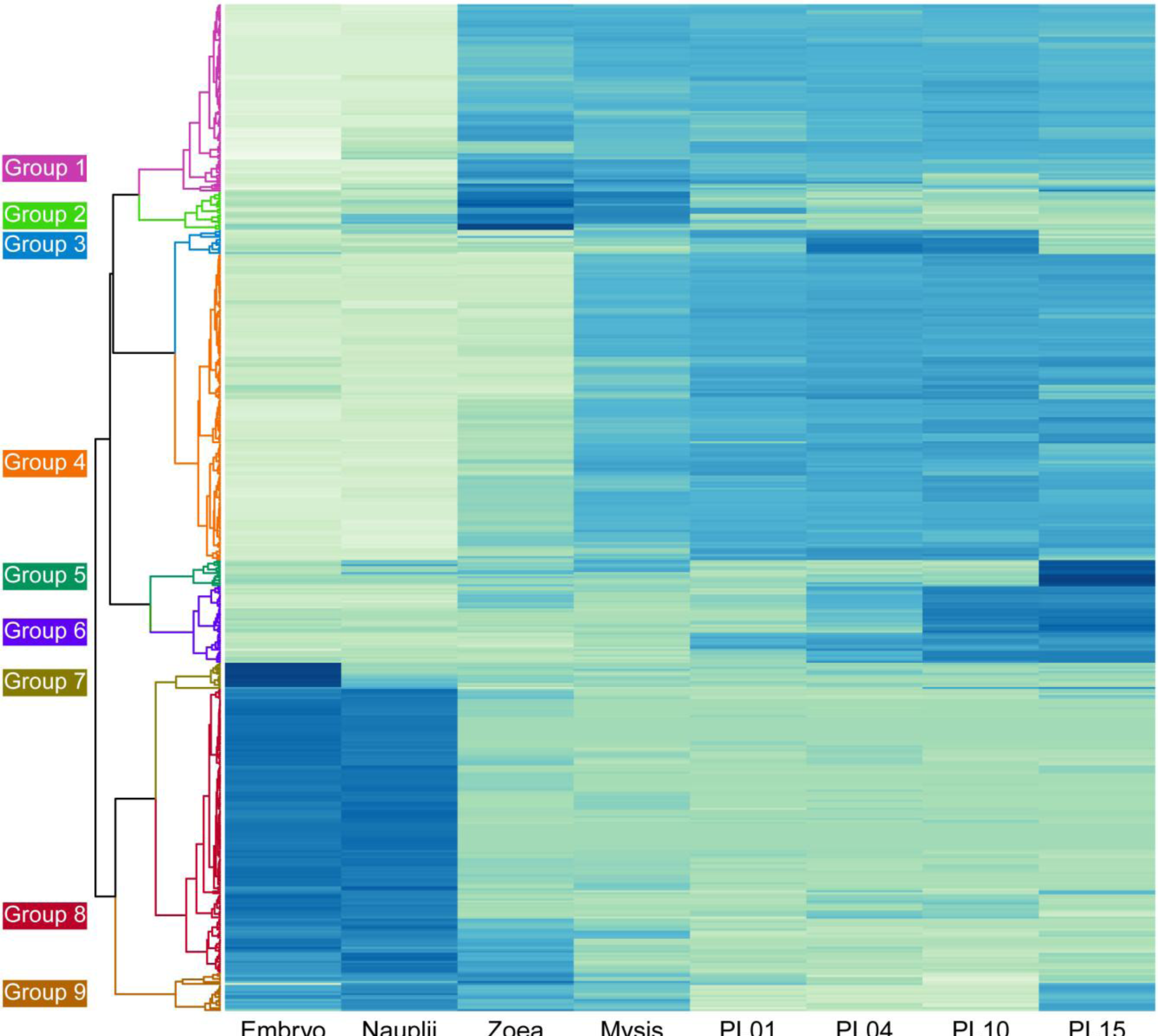
Heatmap and hierarchical grouping of the top 500 differentially expressed genes in the eight larval and post-larval stages examined. Gene expression patterns in each larval/post-larval stage (row) were grouped into nine expression groups based on Pearson’s correlation.

### Identification of long non-coding RNAs

We used the set of 1,047 complete USCOs as the training set for classification of coding and non-coding transcripts. It was determined that a coding potential of 0.2642 was the appropriate threshold to balance classification specificity and sensitivity. In total 79,656 transcripts were classified as lncRNAs and the remaining 154,893 transcripts were classified as mRNAs.

Comparing the lncRNA annotation with the BLASTx annotation, out of the 236,388 contigs 67,960 were uniquely identified as lncRNA, while 13,535 contigs were annotated both as mRNA and lncRNA. At a cluster level, 12,079 out of 58,768 larval clusters (22.6%) and 23,645 out of the 99,203 tissue clusters (23.8%) were uniquely annotated as lncRNA. Detailed results of the lncRNA analysis can be found in Supplementary Table 4.

### KEGG pathway analysis

Annotated contigs were overlaid onto their respective biological pathways using the Kyoto Encyclopaedia of Genes and Genomes (KEGG) pathways. Genes involved in general eukaryotic cellular processes such as RNA replication (Fig. 6) and basal transcription factor sequences (Fig. 7) were well represented in the *P. monodon* transcriptome. As expected, assignments to KEGG pathways in prokaryotes were rare, as were ribosomal RNA assignments. The various biological processes, metabolism and signalling cascades comprising all 235 KEGG pathways to which transcripts were assigned are detailed in Supplementary Table 5.

**Fig. 6.**
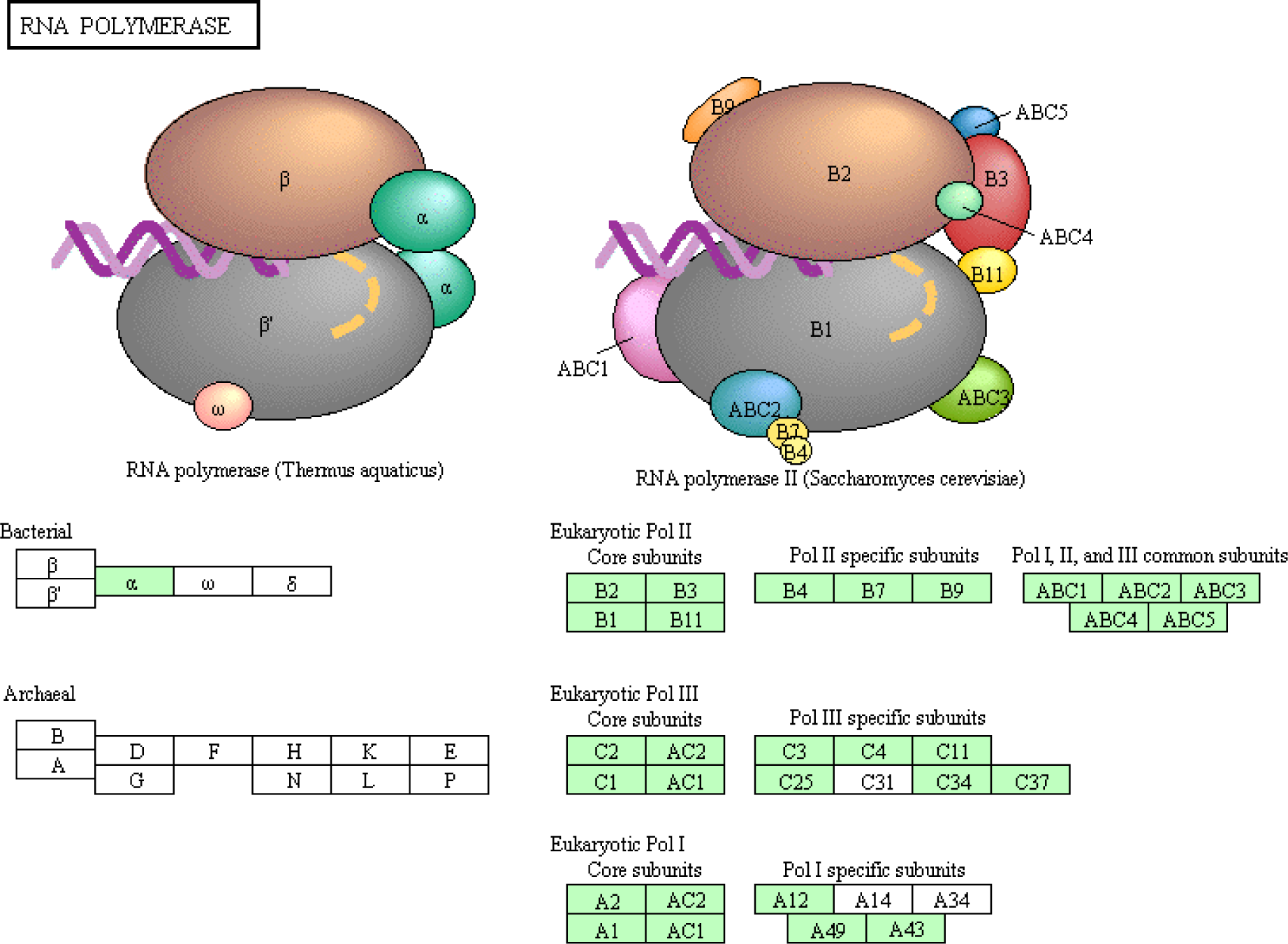
Presence of mRNA contigs that encode for RNA polymerase subunits based on KEGG pathway analysis. Green shading highlights the presence of gene orthologs in the *P. monodon* transcriptome.

**Fig. 7.**
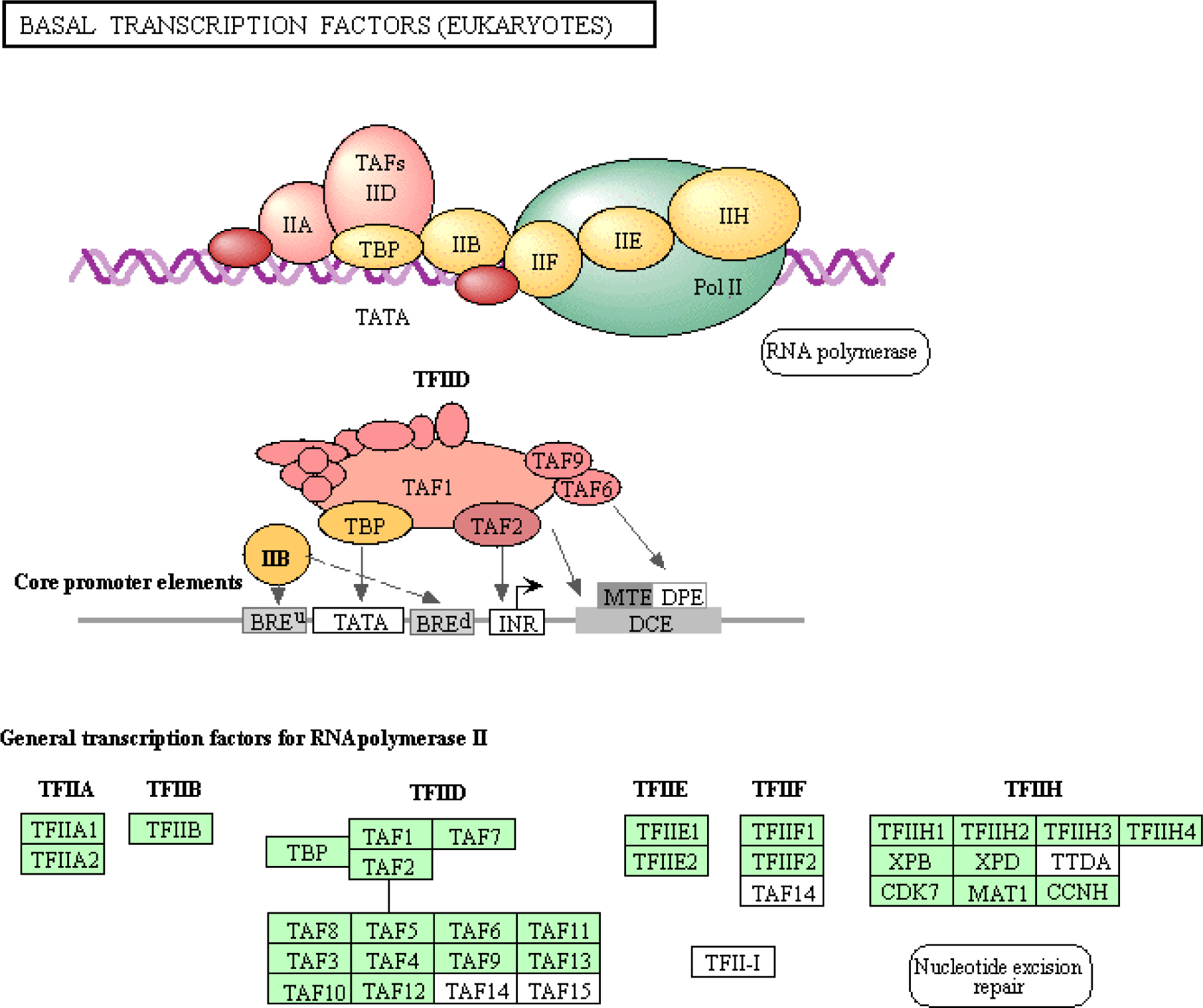
Presence of eukaryotic basal transcription factor sequences based on KEGG pathway analysis. Green shading highlights the presence of gene in the *P. monod*on transcriptome.

### Virus discovery

Interrogating the *P. monodon* transcriptome against the viral subsection of the non-redundant database using BLASTx assigned viral annotations to 12,744 contigs.Detailed information on the viral blast can be found in Supplementary Table 6. Closer inspection of the data identified the vast majority (>99.8%) of these to represent short motifs conserved between eukaryote cell proteins and related homologs viruses with generally large and complex DNA genomes such as giant viruses, poxviruses, herpes viruses and baculoviruses. Additional BLASTx searches of the GenBank nr database using representative contigs confirmed them to be or likely be endogenous shrimp gene transcripts. The remaining 21 contigs had Top Hit E-value scores identifying them to be related most closely to strains of Gill-associated virus (GAV; 4 contigs, longest 26,235 nt), *Penaeus chinensis* hepandenovirus (*Pchi*HDV; 4 contigs, longest 1,884 nt), Wenzhou shrimp virus 2 (WSV2; RdRp, hypothetical protein and G protein contigs, longest 6,891 nt), Deformed wing virus (DWV; 10,133 nt), Wenzhou shrimp virus 8 (WSV8; 6 contigs,longest 4,579 nt), Beihai picorna-like virus 2 (5,277 nt), Wenzhou picorna-like virus 23 (551 nt) and Moloney murine leukaemia virus Pr180 sequence (Mo-MuLV; 2,431 nt). Lastly, over 1200 contigs with homology to phages were detected, some of which related to phage tail protein and tetracycline resistance.

## Discussion

Here we report a comprehensive black tiger shrimp (*Penaeus monodon*) transcriptome assembled from nine tissues, four larval stages and four post-larval stages. The transcriptome was generated to expand the genetic resources available for this species to help investigate the genetic basis behind larval developmental stage transitions and tissue functioning, as well as traits with potential to be exploited commercially for the aquaculture of this and other shrimp species. The aim was therefore to generate a highly complete *P. monodon* transcriptome at the risk of it containing higher levels of transcript redundancy. This was confirmed by BUSCO results which demonstrated the transcriptome to be highly complete (C: 98.2%) with low fragmentation (F: 0.8%) or missing (M: 1.0%) genes but high levels of duplication (D: 51.3%). These assembly statistics are comparable to those obtained by a transcriptome assembly from *L. vannamei*^15^ (C: 98.0%, F: 0.7%, M: 1.3%, D: 25.5%), but greatly exceeded those of another *P. monodon* assembly focussing on gonadial tissue recently made available publicly^10^ (C: 33.7%, F: 44.9%, M: 21.4%, D: 6.8%).As other recent NGS analyses of *P. monodon* have focussed on only one or two tissue types without including any larval stages or biological replicates, generated fewer total reads, or experienced data loss due to quality trimming of low quality reads or low mapping efficiencies^8–11^, these are likely to have missed many transcripts. In contrast, the sequencing and assembly strategy used here covered more tissue types at greater read depth and employed multiple *de novo* assembly tools to reduce assembler bias.

### Functional annotation and comparative analysis

Using the nrA database, 30.0% of transcript clusters found in the nine tissue types and 38.1% of transcript clusters found in the eight larval/post-larval stages analysed were successfully annotated. These annotation levels were comparable to those reported to date in similar studies on different crustaceans^8,15,24,50^. While transcript cluster annotation levels were lower using the SWISS-PROT database compared to the nrA database, the percentage of successful GO-term assignments was substantially higher. In addition to the annotations, analyses were undertaken to identify transcript clusters expressed differentially across tissue types or early life-history stages, irrespective of successful annotation. The identification was done to help provide initial evidence for transcript roles in specific tissue functions or developmental transitions. Despite all efforts made here to improve transcript annotation levels for *P. monodon*, our data reaffirms the need for dedicated functional studies to assign or confirm gene functions of both annotated and unannotated transcript clusters of non-model (crustacean) species.

To our best knowledge, to date only two Penaeid shrimp transcriptome assemblies have been made publicly available^10,15^, restricting comparative analyses of these transcriptomes. A reciprocal MegaBLAST identified 96.8% of the most recent *P. monodon* assembly ^10^ within the transcriptome described here, but only 40.0% of our assembly was found in the earlier assembly. These comparisons confirm that our transcriptome assembly contains many high quality *P. monodon* transcripts not discovered previously.

When compared across species, a reciprocal MegaBLAST showed that the transcriptomes of *P. monodon* (present) and *L. vannamei*^15^ shared approximately 48% of contigs. Since the assembly metrics of the *L. vannamei* transcriptome were similar to those of our *P. monodon* transcriptome, the low number of shared contigs could stem from considerable differences in transcript type or sequence composition between the two shrimp species. As comprehensive comparisons across crustacean species is currently impractical due to restrictions on publicly-available transcriptome assemblies, the potential value of this warrants effort to consolidate transcriptomic data and to establish both centralized and species-specific databases.

### Tissue specific expression

Read count data identified independent clusters of transcripts expressed uniquely within different tissues and clusters that formed distinct groups based on their tissue-specific expression patterns. An important consideration for this type of analysis is the normalized read count cutoff value for each cluster to be considered “unique”, which was arbitrarily set at above 10 in a specific tissue and < 10 in all others. At >100 normalized read counts, only approximately half of the assigned unique clusters were retained, indicating that the expression levels of many of these potentially tissue-specific clusters was relatively low. Among the annotated transcript clusters most highly expressed in female gonad tissue were FAMeT, PEPCK, GPx and nasrat. Functional roles these proteins may play range from the shrimp moult cycle and reproduction^51^, the primary step of gluconeogenesis^52^, preventing oxidative stress^52^, to specifying terminal regions of the embryo^53^. Among the annotated genes expressed most highly in eyestalk tissue was hyperglycaemic hormone (CHH), a key neuropeptide hormone that regulates blood sugar, moulting and reproduction^54^. A subset of transcript clusters highly expressed in lymphoid organ tissue was also highly expressed in gill tissue, most likely due to high concentrations of haemocytes within both tissue types. The majority of genes expressed most highly in hepatopancreas were annotated, potentially reflecting the shared metabolic functions of this organ with those of other animals. Also of much interest were the non-annotated transcripts expressed uniquely in specific tissue types. For example, transcript clusters expressed highly in male gonad were poorly annotated by both databases and included a large proportion of clusters, annotated or not, expressed exclusively in adult tissue types, indicating that male reproductive organs utilize many genes that remain poorly characterized. The grouping of genes with similar expression patterns broadly categorized these transcript clusters into potential functional groups within each tissue type, thereby guiding the selection for more targeted molecular function analyses.

### Larval and post-larval development

Based solely on gene expression patterns, the transcriptome data identified unique groups of transcripts involved in transitions between *P. monodon* early life-history stages. There was a major disparity between the annotation success of transcript groups upregulated in early or late stage embryogenesis, highlighting how poorly early developmental pathways have been characterized in crustaceans. Also of significance was the presence of orthologs of the *Spaetzle* gene, known in *Drosophila* flies to establish the dorso-ventral patterning of the early embryo^55^ among transcript clusters detected consistently across later larval and post-larval stages.Since each larval and post-larval stage sequenced comprised a pool of several hundred individuals, quantitative and/or spatial transcript expression patterns would be required to draw further functional conclusions. Nevertheless, the data reported here will benefit from similar data on other shrimp and crustacean species, particularly for transcript clusters expressed exclusively in embryo with no significant homology to currently known genes.

### Identification of long non-coding RNAs

Long non-coding RNAs (lncRNA) are a type of transcript that have many common features with traditional coding mRNA, including 5’ capping, splicing and 3’ polyadenylation^56–58^. The nature of lncRNAs is still poorly understood, and it is likely that lncRNAs are in fact a heterogeneous group of transcripts with regulatory functions that are not actively translated into proteins^59^. Thus, their main characteristics are the lack of open reading frames (ORFs) or the presence of non-canonical ORFs in the mature transcript. The biological roles of lncRNAs range from regulation of gene expression, and control of translation, to imprinting. As such, they have been linked to X chromosome inactivation in humans^60^, genomic imprinting^61^ and cancer^62,63^.

Due to the lack of a known lncRNA database in shrimp that can be used for their identification, we used FEELnc which scores each transcript according to its coding potential and then selects a threshold score to classify the transcripts into coding or non-coding^45^. This software is particularly useful for non-model species because in the absence of an lncRNA training set, it generates a simulated training set using debris from high confidence coding transcripts. In fly data, this approach showed an MCC value of 0.754 with an accuracy of 0.868^45^.

In this study, 79,656 transcripts were classified as lncRNAs, of which 67,960 (85.3%) could not be aligned to any protein database. As expected, the use of a non-model organism and the lack of a set with known lncRNA for training led to the ambiguous classification of 13,535 transcripts with low protein-coding potential but clear alignments to known proteins in curated databases. Classification of these transcripts is the first step towards understanding their roles in the development and regulation of gene expression in *Penaeus monodon*.

### KEGG pathways

Annotated transcript clusters mapped into 235 KEGG pathways (Supplementary Table 3), which have been broadly classified into functional groupings such as general metabolism (e.g. TCA cycle, xenobiotic metabolism, immunity, reproduction), nutritional metabolism (e.g. proteins, lipids, carbohydrates, vitamins), cellular processes (e.g. DNA replication, protein trafficking, apoptosis), biological processes (e.g. circadian rhythm, olfaction and taste, digestion and absorption) and signalling pathways (e.g. PI3K-Akt, MAPK, axis formation, TGF-beta). In general, core pathways such as citrate cycle, oxidative phosphorylation, ribosome biogenesis and RNA/DNA polymerases were better represented than more specific pathways such as the pentose and glucuronate interconversion pathway, or the ascorbate and aldarate metabolism pathway. Furthermore, arthropod specific pathways were generally better represented. For example, the general circadian rhythm pathway was missing several homologs, while the fly specific circadian rhythm pathway was complete. This could be explained by transcripts not sharing sufficient homology with the known genes used for the KEGG analysis and therefore failing to be annotated. Particularly for those pathways highly-conserved among other eukaryotes, the existence of unique transcripts suggests that Penaeid shrimp and possibly crustaceans in general might use metabolic mechanisms differing from eukaryote species studied to date. Their existence also highlights the need for high-quality genome assemblies for shrimp and other crustacean species, overlaid with isoform, tissue-specific and developmental stage transcript expression data, to either help predict gene functions or direct gene knockdown studies, using RNA interference processes as an example, to empirically ascribe functions to novel genes.

### Virus discovery

Several RNA transcripts and/or genome sequences likely to be from viruses were discovered in the *P. monodon* transcriptome. This was not unexpected considering that it was generated from multiple individuals, tissue types and larval/post-larval stages, as shrimp are co-infected commonly with multiple viruses and as there are several viruses known to be endemic in *P. monodon* populations indigenous to different regions of Australia^64–67^. The presence of near full-length ssRNA genome sequences for viruses such as gill-associated virus (GAV, 26,235 nt) and white spot virus 2 (WSV2, 10,542 nt) provided additional validation of the methods used to synthesize and assemble the transcriptome, and to its completeness as demonstrated by various metrics measuring the nature and number of endogenous gene transcripts. The detection of a ssDNA virus, hepandenovirus, within the transcriptome, presumably detected in a replicative phase, indicates the application of this technique as a tool to also detect the presence of viruses with DNA genomes.

In addition to known endemic viruses, the transcriptome contained full-length or near full-length RNA transcripts related closely to the recently-described shrimp viruses WSV2 and WSV8^68,69^ unknown until now to occur in Australian *P. monodon*.Moreover, it contained a long transcript (10,133 nt) 95.0% identical to the full-length ssRNA genome of deformed wing virus (DWV), a virus of Varroa mites that is transmitted to honeybees^70^, and one of a rapidly expanding number of *Iflavirus* species now being discovered in diverse insect species also including beetles, wasps, caterpillars and moths^71^. As essentially all DWV-like genome sequence reads in this study originated from the stomach of a single individual shrimp, they were potentially derived from a virus-infected honeybee or mite-infested honeybee ingesting by this shrimp. While honeybees infested with Varroa mites have been detected recently in North Queensland not far from where the shrimp was collected^72^, DWV itself has not been detected in a comprehensive recent study^73^.The present study therefore represents the first detection of a DWV-like genome in Australia, although the origin remains unknown. This reinforces both the strength of the technology in detecting unknown pathogens and also the potential difficulty in interpretation of transcriptome results.

A couple of long transcripts of suspected viral origin and expressed across multiple tissue types were also identified. One of these possessed significant BLASTx homology to the reverse transcriptase (RT)-like component of hypothetical protein 1 of Beihai picorna-like virus 116 discovered recently in blue swimmer crabs (*Portunus pelagicus*)^69^. The other possessed substantial homology to the RT component of the Mo-MuLV Pr180 polyprotein and was expressed across all tissue types except the lymphoid organ, suggesting it to be from a mobile element such as a poly(A)-type retrotransposon or retrovirus^74^. However, determining whether these transcripts containing RT sequences are viral in origin, or represent the products of endogenous retrotransposons like others now being reported in shrimp^75^ will require further investigation, as will the nature of the strains, host and distribution ranges,prevalence and potential pathogenicity of the new viruses discovered in the transcriptome.

### Conclusions

This study describes the assembly of a comprehensive and high quality transcriptome from nine different tissue types, and eight larval and post-larval early life-history stages of the black tiger shrimp, *Penaeus monodon*. It also summarizes the number and nature of specific transcript clusters differentially expressed in different tissue types and larval stages, and the Clusters were functionally annotated and mapped to 235 KEGG pathways. Unique transcript clusters and cluster groups were defined across distinct tissues and early life-history stages, providing initial evidence for their roles in specific tissue functions or developmental transitions. The current transcriptome provides a valuable resource for further investigation of directing gene-function studies to increase basic functional biology knowledge in shrimp and for investigating molecular basis of traits of relevance to the aquaculture of shrimp. While the current transcriptome already provides an improved resource for *P. monodon*, further effort is required using long-read sequencing data, such as provided by PacBio, to better resolve genes at isoform level. Lastly, this high-quality *de novo* assembly and data set are publically available and will hopefully support research projects that underpin transformational advances in how we culture shrimp globally.

## Declarations

### Availability of data and material

Raw data, assembly and bioinformatics scripts will be made freely available online upon publication.

### Authors’ contributions

RH: conceptualised, developed and oversaw the project, performed sampling and carried out RNA extractions, developed and performed the transcriptome assembly, quality assessment and differential gene analysis, and wrote the manuscript. NMW: conceptualised and developed the project, performed sampling, led components of data analysis and interpretation and wrote the manuscript. LG: developed the transcriptome assembly bioinformatics pipeline. JDM: carried out the lncRNA analysis and assisted with the transcriptome assembly bioinformatics pipeline. JG: carried out sampling and RNA extractions, and assisted with development of the transcriptome assembly bioinformatics pipeline, and reviewed the manuscript. SM: assisted with the bioinformatic analysis of the differential gene expression data and reviewed manuscript. MT: oversaw the library preparation and sequencing, and reviewed the manuscript. KS: conceptualised and developed the project and reviewed manuscript. EG: reared and sampled the larval stages. DD: conceptualised and developed the project. Coordinated facilities and resources for larval and adult prawn production and reviewed manuscript. MS: provided prawn tissue samples, conceptualised and developed the project, assisted with data interpretation and reviewed manuscript. JC: assisted with the interpretation and writing of the viral analysis, and edited manuscript. KC: assisted with the interpretation and writing of the viral analysis, and reviewed manuscript. GC: conceptualised and developed the project and reviewed manuscript. MK: conceptualised and developed the project and reviewed manuscript. HR: conceptualised and developed the project and reviewed manuscript. GM: conceptualised and developed the project, performed sampling, provided advice on sequencing strategies and data interpretation and reviewed manuscript. KRZ: conceptualised and developed the project and reviewed manuscript. DRJ: conceptualised and developed the project, oversaw coordination of project activities and reviewed manuscript

All authors read and approved the final manuscript.

## Acknowledgements

We thank Andrew Foote, Gopala Krishna, Sarah Berry and Tansyn Noble for assistance in organizing and collecting tissue samples. Research Funding for this project was from the Australian Research Council Industrial Transformation Research Program IH130200013.

## Supplementary Material

**Supplementary Table 1** contains all annotation results from the blast against SwissProt and nrA (arthropod subsection of the nr database), including blast metrics, GO terms, interpro scan results and simplified lncRNA results.

**Supplementary Table 2** contains the top 2,000 differentially expressed genes in the nine different tissue types used in the heatmap. The table shows normalised expression values for each sample, the nine groupings based on Pearson’s correlation presented in the heatmap, and the associated SwissProt and nrA annotations.

**Supplementary Table 3** contains the top 500 differentially expressed genes in the eight early life-history stages used in the heatmap. The table shows normalised expression values for each sample, the nine groupings based on Pearson’s correlation presented in the heatmap, and the associated SwissProt and nrA annotations.

**Supplementary Table 4** contains the detailed results of the lncRNA analysis using the FEELnc pipeline.

**Supplementary Table 5** contains links to the 235 KEGG pathway figures based on the transcriptome generated in this study.

**Supplementary Table 6** contains all successful blast hits against the viral subsection of the nr database.

